# SAVED by a toxin: Structure and function of the CRISPR Lon protease

**DOI:** 10.1101/2021.12.06.471393

**Authors:** Christophe Rouillon, Niels Schneberger, Haotian Chi, Martin F. Peter, Matthias Geyer, Wolfgang Boenigk, Reinhard Seifert, Malcolm F. White, Gregor Hagelueken

## Abstract

CRISPR antiviral defense systems such as the well-known DNA-targeting Cas9- and the more complex RNA-targeting type III systems are widespread in bacteria and archea ^1, 2^. The type III systems can orchestrate a complex antiviral response that is initiated by the synthesis of cyclic oligoadenylates (cOAs) upon foreign RNA recognition ^3–5^. These second messenger molecules bind to the CARF (CRISPR associated Rossmann-fold) domains of dedicated effector proteins that are often DNAses, RNAses, or putative transcription factors ^6^. The activated effectors interfere with cellular pathways of the host, inducing cell death or a dormant state of the cell that is better suited to avoid propagation of the viral attack ^7, 8^. Among a large set of proteins that were predicted to be linked to the type III systems ^9, 10^, the CRISPR-Lon protein caught our attention. The protein was predicted to be an integral membrane protein containing a SAVED-instead of a CARF-domain as well as a Lon protease effector domain. Here, we report the crystal structure of CRISPR-Lon. The protein is a soluble monomer and indeed contains a SAVED domain that accommodates cA_4_. Further, we show that CRISPR-Lon forms a stable complex with the 34 kDa CRISPR-T protein. Upon activation by cA_4_, CRISPR-Lon specifically cleaves CRISRP-T, releasing CRISPR-T_23_, a 23 kDa fragment that is structurally very similar to MazF toxins and is likely a sequence specific nuclease. Our results describe the first cOA activated proteolytic enzyme and provide the first example of a SAVED domain connected to a type III CRISPR defense system. The use of a protease as a means to unleash a fast response against a threat has intriguing parallels to eukaryotic innate immunity.

## Main

CRISPR (Clustered Regularly Interspaced Short Palindromic Repeats) is a bacterial and archaeal adaptative immune system that enables microorganisms to fend off attacks by mobile genetic elements such as phages, viruses, or plasmids ^11^. The protein complex Cas1-Cas2 captures short DNAs from invaders and integrates them as “memories” into a CRISPR locus ^12^. Transcripts of these “memories” are processed into small CRISPR RNAs (crRNAs) and integrated into large ribonucleoprotein (RNP) complexes, which can sense the presence of a matching foreign nucleic acid in the cell ^13^. Once a foreign nucleotide sequence is detected, an antiviral response is triggered. Depending on the type of CRISPR system ^1^, this response can be markedly different, ranging from simple cleavage of the invading nucleic acid by the RNP as in the case of Cas9 ^14^, to a complex multipronged defense strategy as found in type III CRISPR systems ^6^. For the latter, the Cas10 subunit of the RNP has a cyclase activity that converts ATP into a recently discovered class of cyclic oligoadenylates (cOAs) upon viral RNA recognition ^3–5^. The cOAs are constructed from 3 to 6, 3’-5’ linked AMP units ^15^ and act as second messengers by binding to proteins harboring a CARF (CRISPR-associated Rossmann-fold) domain ^16^. There is a large variety of CARF proteins linked to effector domains with functions ranging from RNA cleavage, supercoiled DNA nicking, dsDNA cleavage to transcription modulation ^6, 17–22^. The downstream effects of those cOA-activated proteins can lead to an abortive infection or a dormant state of the cell, enabling it to weather the phage attack ^7, 8^.

Recently, two bioinformatic teams attempted to catalog CARF-domain containing proteins that are likely linked to a functional type III system in bacterial and archaeal genomes ^9, 10^. Together, the studies revealed more than 100 such proteins, including several membrane proteins and many proteins with currently unknown functions. Another study showed that some of those type III-associated proteins contain a SAVED domain (‘SMODS-associated and fused to various effectors domains’; SMODS being the acronym for ‘second messenger oligonucleotide or dinucleotide synthetase’ ^23^) instead of CARF, reminiscent of the recently discovered CBASS system (‘cyclic-oligonucleotide-based antiphage signaling systems, ^24^). One of these proteins is CRISPR-Lon (originally termed Lon-CARF), a 60 kDa protein with two predicted transmembrane helices, a Lon-protease domain and a SAVED4 domain ^25^.

Here we report the high-resolution crystal structure of the CRISPR-Lon protease from the thermophilic bacterium *Sulfurihydrogenibium sp.* YO3AOP1. We find that CRISPR-Lon is a soluble monomer and forms a complex with CRISPR-T, a small 34 kDa protein encoded by an adjacent gene in the locus. Once activated by cA_4_, CRISPR-Lon proteolytically cleaves CRISPR-T, releasing CRISPR-T_23_, a 23 kDa fragment predicted to have a MazF-like ^26, 27^ RNAse activity.

### Structure of CRISPR-Lon

A synthetic, codon-optimized variant of the CRISPR-Lon gene from *Sulfurihydrogenibium sp.* YO3AOP1 (UNIPROT-ID: B2V8L9) was cloned into the pET11a vector and expressed in *E. coli.* Although predicted to include two trans-membrane helices ^9, 10^, we noticed that the protein was found in the soluble fraction of the cell lysate and behaved as a stable monomer during size exclusion chromatography (Supplemental Figure 1 and below). The pure protein readily crystallized, but several rounds of optimization were necessary to obtain well diffracting crystals. No suitable structural model was available at the time, and we prepared a seleno-methionine derivative, which crystallized under the same conditions. The structure of CRISPR-Lon was solved at 2.1 Å by single-wavelength anomalous dispersion phasing and refined to R/R_free_ values of 19.3/22.5. The electron density was well defined and the model had good geometric parameters (Supplemental Table 1, Supplemental Figure 2) ^28, 29^. As expected from gel filtration experiments (Supplemental figure 1 and below), the crystal packing does not indicate any stable multimeric forms of CRISPR-Lon (Figure 1A) ^30^.

**Figure 1:**
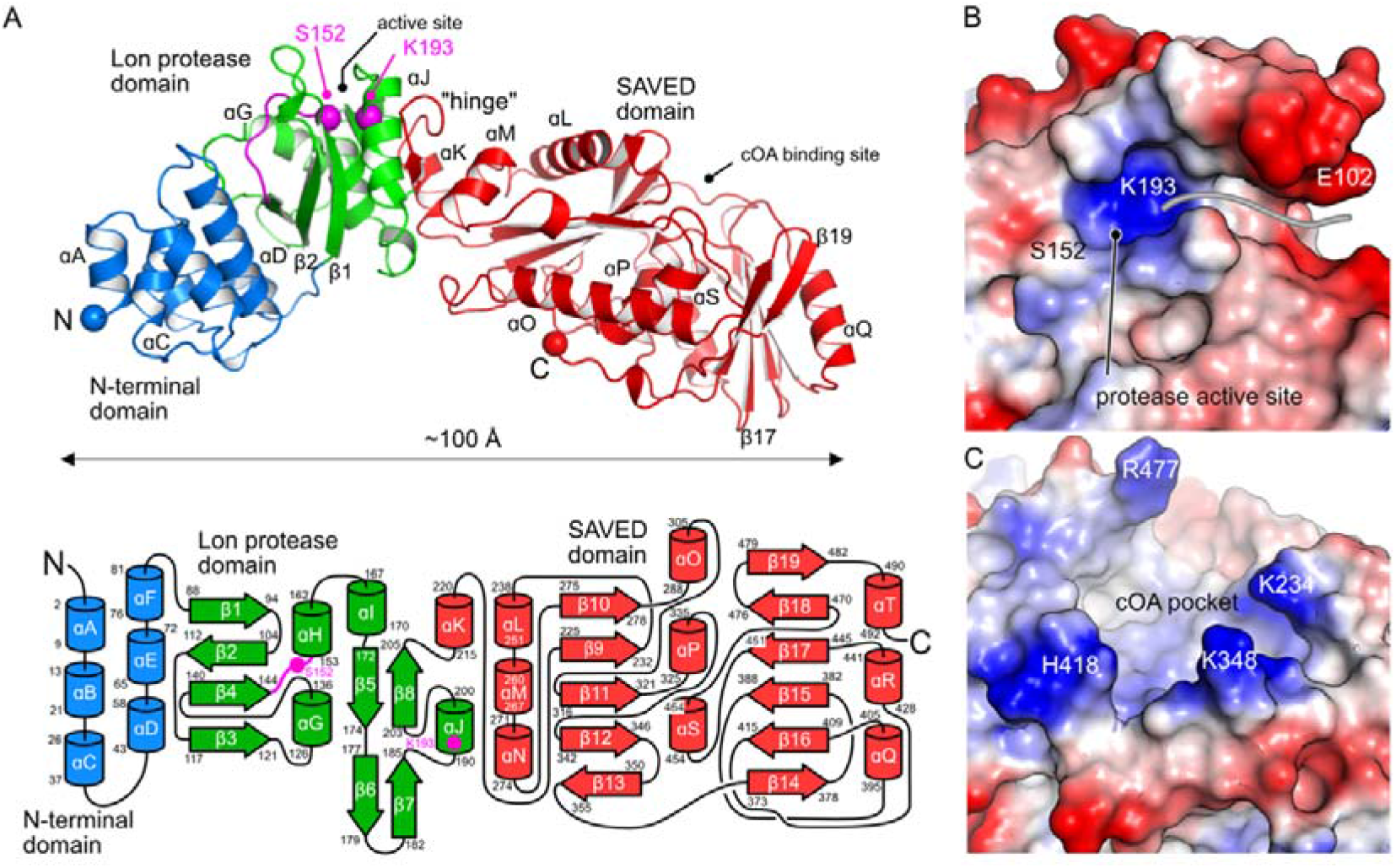
Structure of CRISPR-Lon. **A)** Overall structure and topology of CRISPR-Lon. The structure is shown as a cartoon model and the individual domains are color-coded as in the topology diagram (bottom). The N- and C-termini, as well as the protease active site residues, are marked by spheres. The positions of key structural elements are indicated. **B)** The entrance to the Lon protease active site. The electrostatic surface of the protein is shown with red representing negatively charged and blue representing positively charged areas. The position of the catalytic S152-K193 dyad is indicated and the tentative position of a substrate peptide is indicated by a grey line. **C)** The cOA binding site with surface electrostatics (blue - positive, red – negative). For orientation, selected amino acid residues are labeled.

The Lon-protease domain consists of a four-stranded mixed β-sheet (β 1-4), sandwiched between □D of the N-terminal domain and □G, H, J on the other side. It is structurally related to the ATP-dependent Lon protease from *Thermococcus onnorineus* (PDB-ID: 3K1J, DALI Z-score: 12.8, Supplemental Figure 3,^31^), the ATP-independent VP-4 protease from Tellina virus (3P06, Z-score: 11.0, ^32^), and the RadA helicase protein from *Streptococcus pneumoniae*, which lacks the protease active site residues (PDB-ID: 5LKM, Z-score 11.7, ^33^). In CRISPR-Lon, the catalytic Ser-Lys dyad, a hallmark of Lon proteases, is formed by S152 (loop β4-□H) and K193 (□J)(Figure 1A). As in other Lon protease structures, a sulfate ion from the crystallization condition binds to the active site entrance (Supplemental Figure 2). The protease active site lies at the end of a narrow channel that presumably binds the substrate peptide (Figure 1B). Supplemental Figure 4A shows a superposition of the Lon-protease domain of CRISPR-Lon with the acyl-enzyme intermediate state of the yellowfin ascites virus ATP-independent Lon protease (PDB-ID: 4IZJ, Z-score: 9.8, ^34^). In that structure, the substrate peptide attaches to the central β-sheet as a fifth strand. The corresponding region of CRISPR-Lon, loop β1/ β 2, is slightly disordered and will likely adopt a similar conformation in the substrate-bound form. The superposition with 4 IZJ allowed us to deduce the position of the P1 site in the protease active site of CRISPR-Lon. As illustrated in Supplemental Figure 4A, only peptides with small side chains such as Ala or Gly would fit into this site.

The C-terminal part of CRISPR-Lon folds into a SAVED4 domain ^25^, consisting of two pseudo-symmetric CARF-like domains with a pseudo-two-fold axis running between helices □P and □S (Figure 1A). Interestingly, the TMHMM 2.0 server predicted that the latter two helices and the directly preceding β-strands form TM helices or at least membrane associated helices ^9, 35^ (Figure 1, Supplemental Figure 2B). The SAVED domain forms an extensive cavity on its molecular surface (Figure 1C). Several positively charged residues at the entrance to this pocket presumably bind the phosphate groups of the cOA ligand. By molecular modelling, we found that of cA_3-6_, cA_4_ showed the best steric fit into this pocket (Supplemental Figure 5A). The cOA binding domains of the Cap4, Can1 and Cap5 proteins were identified as close structural homologs using the DALI server (PDB-ID Cap4: 6VM6, Z-score: 17.1, ^24^; PDB-ID Cap5: 7RWK, Z-score: 9.9 ^36^; PDB-ID Can1: 6SCE, Z-score 8.0, ^19^).

The superpositions in Supplemental Figure 5 show that despite the low sequence identities (Cap4: 14 %, Can1: 9 %, Cap5: 13 %), the fold of the CARF-like domains is conserved. Interestingly, the position of the effector domain relative to the SAVED- or CARF domains is entirely different between the four structures (Supplemental Figure 5BC). Comparing the cOA binding sites of Cap4 and CRISPR-Lon shows that the surface loops of the SAVED domains lead to a very different shape of the cOA binding site (Supplemental Figure 5AB).

The N-terminal domain of CRISPR-Lon forms a bundle of six □-helices (□A-F). The DALI server detected structural similarities (Z-score 6.2) to p60-N, the N-terminal protein-protein interaction domain of Katanin in the p60p80-CAMSAP complex (PDB-ID: 5OW5, ^37^). We noted a surface cluster of hydrophobic residues (W28, L6, V14, L18) formed by □A-C as a possible candidate for a protein-protein interaction interface.

### CRISPR-Lon is activated by cA_4_ and specifically cleaves a putative MazF-like toxin

We used the WebFLAGs server ^38^ to study the gene neighborhood of CRISPR-Lon homologs and noticed a small 812 bp open reading frame (271 amino acids, 32 kDa, UNIPROT-ID: B2V8L8) with no annotated function upstream of the CRISPR-Lon gene (Supplemental Figure 6). We analyzed its sequence with HHPRED ^39^ and found homologies to the MazF toxin in the N-terminal half of the protein and weak homologies to DUF2080, a “domain of unknown function” in the C-terminal half (Figure 2A, Supplemental Figure 7A). We predicted the structure with the deepmind/alphafold2 algorithm ^40^. The software produced a model of a two-domain protein with a ∼23 kDa- and a ∼10 kDa domain connected by an apparently flexible loop (Figure 2B). The structural model was submitted to the DALI server ^41^, revealing structural similarities to MazF-like toxins (Figure 2AB) and various immunoglobulin fold containing proteins (C-terminal fragment, Figure 2AB). Interestingly, the predicted structure appears as a structural mimic of the MazF/E complex with helices □A, □D, and □E blocking the region that binds to the ssRNA target in MazF in a similar fashion to MazE (Supplemental Figure 7C ^42^). Helix □ D of the predicted structure has a large number of arginine and lysine sidechains pointing towards the solvent, suggestive of a nucleic acid interaction interface. We decided to investigate, whether this protein (named “CRISPR-T” below) is the target of the CRISPR-Lon protease. We overexpressed the CRISPR-T gene in *E. coli* and purified it to near homogeneity (Supplemental Figure 8). CRISPR-Lon, CRISPR-T, and different cOAs (3, 4, 5, 6) were mixed at 1:1:1.5 molar ratios and incubated at 60 °C for one hour.

**Figure 2:**
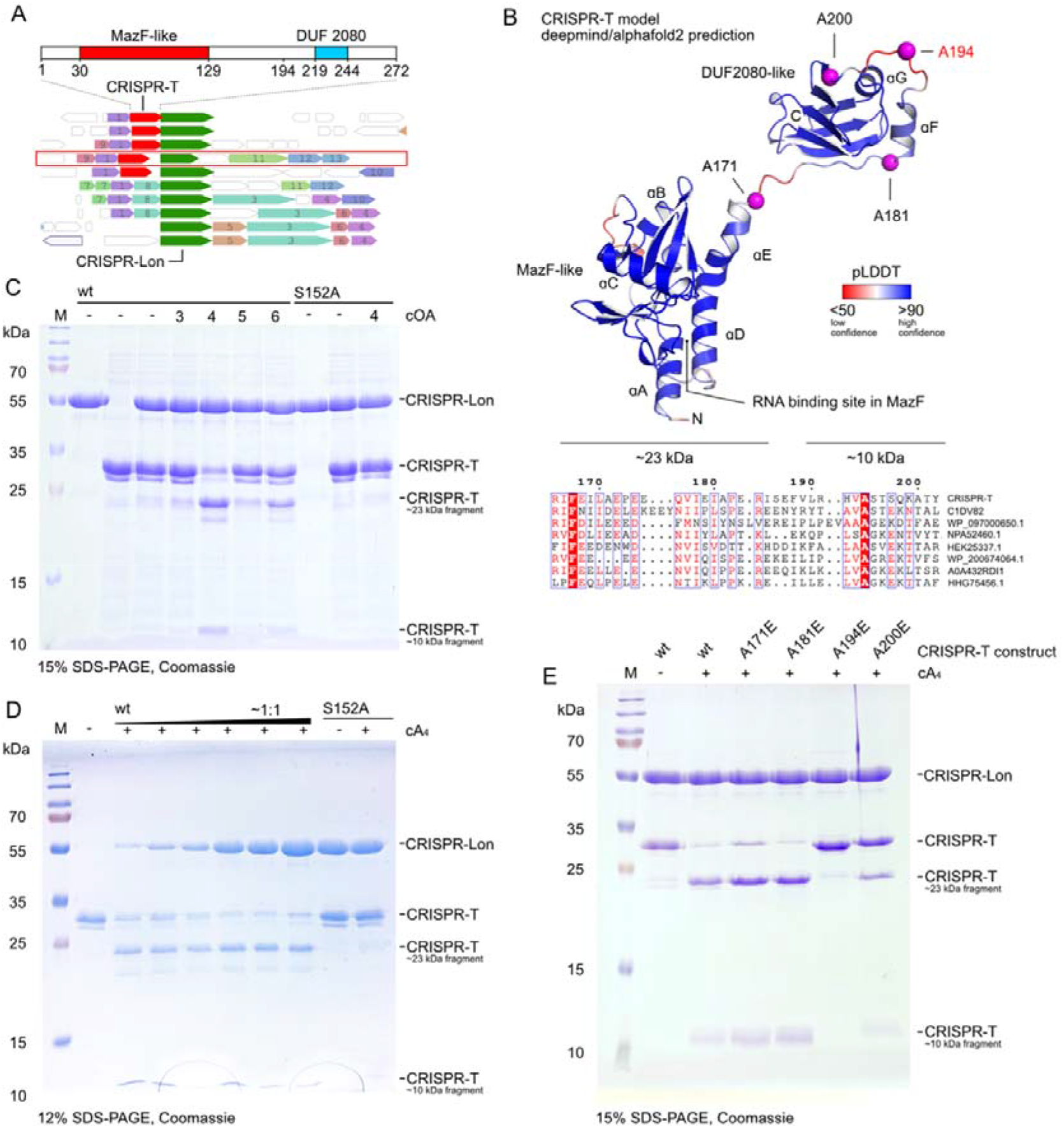
CRISPR-Lon is activated by cA4 and cleaves CRISPR-T. **A)** The WebFLAGs server ^38^ was used to investigate the genomic neighborhood of CRISPR-Lon (green). The primary structure of CRISPR-T (red) is shown on top. Regions with homologies found by HH-PRED ^39^ are marked. **B)** A structural prediction (deepmind/alphafold2, ^40^) of CRISPR-T. The protein is shown as cartoon and colored according to the prediction confidence. The alignment on the bottom ^58^ shows that the identified P1 site at A194 is conserved among CRISPR-T homologs. **C)** SDS-PAGE analysis of CRISPR-Lon induced cleavage of CRISPR-T. **D)** Cleavage experiment as shown in C) but using only cA_4_ and varying concentrations of CRISPR-Lon. **E)** Mutational analysis of potential CRISPR-Lon cleavage sites in CRISPR-T.

Strikingly, we found that in the presence of cA_4_, CRISPR-T was indeed cleaved by CRISPR-Lon. SDS-PAGE analysis revealed two distinct cleavage products with molecular weights of ∼23 kDa and ∼10 kDa (Figure 2CD). The activity for the other cOAs was significantly lower but detectable (Figure 2C). We repeated the experiment with an S152A variant of CRISPR-Lon, which lacks the nucleophilic serine needed for the peptidase activity. This variant showed no protease activity, proving that indeed, the CRISPR-Lon protease active site was responsible for the observed proteolytic activity (Figure 2C).

The peptide sequences of the two cleavage fragments were determined with peptide mass fingerprinting (Supplemental Figure 9). This analysis confirmed that the ∼23 kDa (CRISPR-T_23_) fragment corresponds to the N-terminal two-thirds of the CRISPR-T protein and the ∼10 kDa fragment (CRISPR-T_10_) to the C-terminal one-third. Based on this result and considering the predicted structure (Figure 2B), we narrowed down the location of the cleavage site to the stretch of residues between amino acids ∼170-200 of CRISPR-T. As mentioned above, our CRISPR-Lon structure suggested that only peptides containing an alanine or glycine as the P1 residue will likely fit into the active site of CRISPR-Lon. We created glutamic acid mutants of all four alanine residues in the cleavage region: A171, A181, A194, and A200 (Figure 2B, magenta spheres; the stretch of residues does not contain any glycine). Peptidase assays with all four CRISPR-T mutants were conducted and only the A194E mutation abolished the cleavage completely. Around this position, the amino acid sequence is N-_189_VLRHVA**/**ST_196_-C, where A194 is very likely the P1 residue (Figure 2E). Notably, A194 is conserved amongst CRISPR-T homologs (Figure 2B, bottom). The peptide fingerprint data in Supplemental Figure 9 also supports this conclusion, as for CRISPR-T_23_, the peptide coverage extents exactly to the identified cleavage site. Thus, the molecular weights of the CRISPR-T cleavage products are 23.0 kDa (N-terminal fragment, CRISPR-T_23_) and 8.7 kDa (C-terminal fragment, CRISPR-T_10_), fitting to the sizes observed by SDS-PAGE (Figure 2C).

### CRISPR-Lon and CRISPR-T form a stable complex, releasing a MazF-like toxin upon cA_4_ induced cleavage

We wondered whether CRISPR-Lon and CRISPR-T form a stable complex. To test this, we injected the individual proteins and equimolar mixtures onto a Superose 6 10/300 column connected to a SEC-MALS system (Figure 3A). CRISPR-Lon alone eluted in a single peak at 17.1 ml corresponding to an experimental molecular weight (MW_MALS_) of 52.0 kDa, fitting its theoretical molecular weight (MW_theor_) of 57.7 kDa. The CRISPR-T protein eluted at 17.9 ml with an MW_MALS_ of 30.5 kDa, also close to the expected value (MW_theor_ 34.0 kDa). The 1:1 mixture of the two proteins resulted in a single elution peak at 16.2 ml. The MW_MALS_ of the complex was 82.4 kDa, suggestive of a 1:1 complex of the CRISPR-Lon and CRISPR-T proteins (52 kDa + 30.5 kDa).

**Figure 3:**
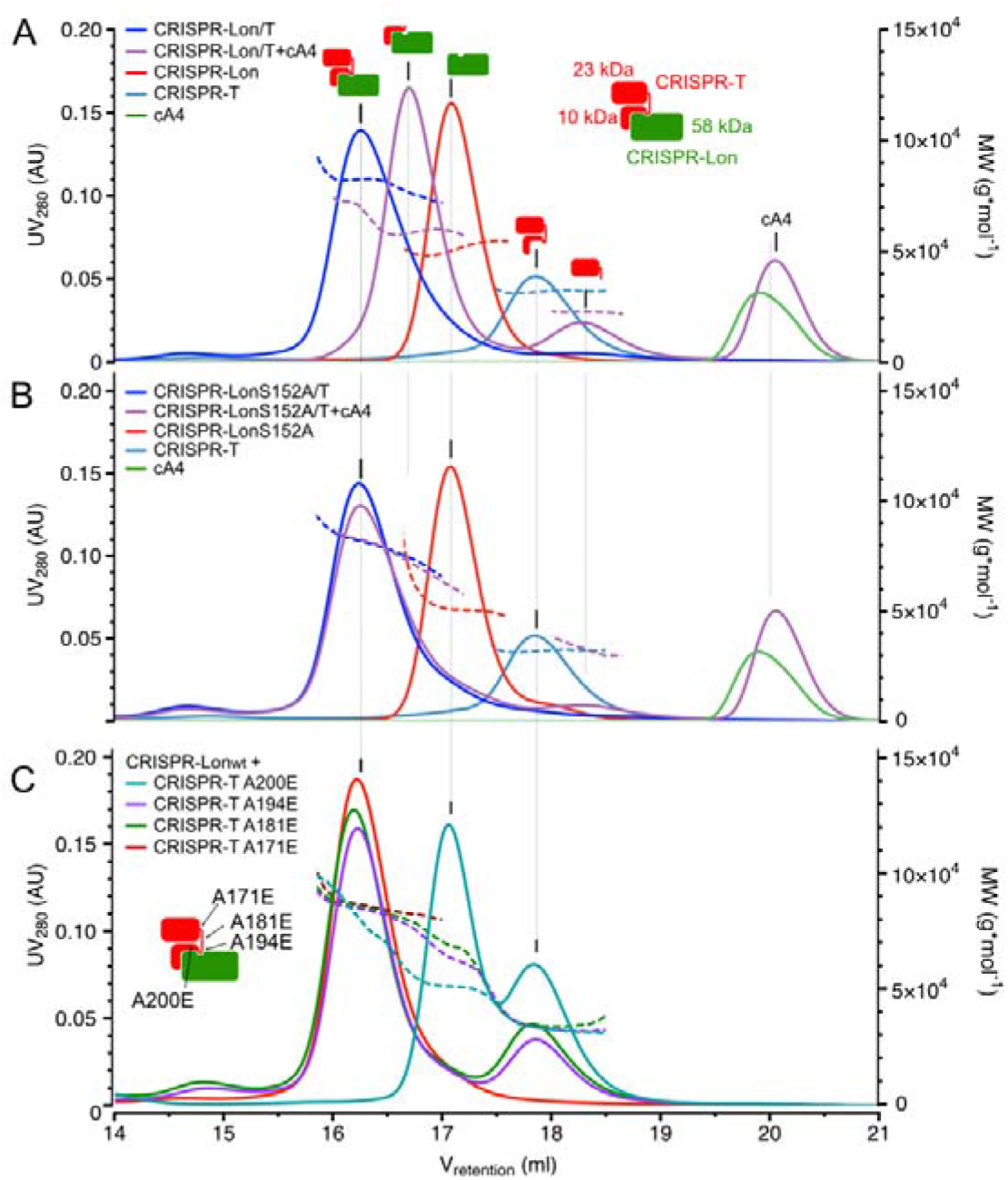
CRISPR-Lon forms a 1:1 complex with CRISPR-T. **A)** SEC-MALS traces (solid lines: UV_280_, dashed lines: MW_MALS_) of proteolysis reactions with different combinations of CRISPR-Lon wt, CRISPR-T wt, and cOA. The schematic indicates the molecular species behind the individual peaks. **B)** Same as A) but with CRISPR-Lon_S152A_ **C)** Binding of CRISPR-Lon wt to the indicated CRISPR-T mutants in the absence of cA_4_. The schematic indicates the position of the mutant in the CRISPR-Lon/T complex.

Next, we repeated the experiment but incubated the CRISPR-Lon/T complex with a slight excess (1:1.1) of cA_4_ (Figure 3B). The SEC-MALS run showed three elution peaks. The first peak at 16.7 ml with an MW_MALS_ of 60.8 kDa nicely agrees with the sum of the MWs of CRISPR-Lon (MW_MALS_ 52.0 kDa) plus CRISPR-T_10_ (MW_theor_ 8.7 kDa). An additional peak at 18.3 ml with an MW_MALS_ of 21.5 kDa fits the size of CRISPR-T_23_ (MW_theor_ 23.0 kDa). The cA_4_ fraction eluted at 20 ml similar to the cA_4_ control and hence did not bind strongly to the complex. We repeated this set of experiments with the inactive CRISPR-Lon S152A variant. Again, the CRISPR-Lon/T complex was observed but the addition of cA_4_ did not lead to the observed split into three elution peaks (Figure 3B). Also here, cA_4_ eluted in a separate peak indicating that the molecule does not bind strongly to the intact CRISPR-Lon/T complex. Finally, we checked whether the four CRISPR-T cleavage-site variants can still form a complex with CRISPR-Lon (Figure 3C). Whereas A171E, A181E, and the P1 site variant A194E still assembled into a 1:1 complex with the protease, the A200E variant did not. A possible explanation would be that A200 is part of the complex interface. The glutamate at this position would then weaken the interaction, leading to the reduced cleavage efficiency observed in Figure 2E.

We proceeded to test for a ribonuclease activity associated with the activated toxin (Supplemental Figure S10). CRISPR-Lon and CRISPR-T were incubated in the presence or absence of activator cA_4_, and full proteolytic digestion of CRISPR-T was observed after 60 min at 60 °C (Supplemental Figure 10A). The cleaved and uncleaved protein mixtures were then incubated with 5 different fluorescent RNA molecules to test a variety of RNA sequences. After 60 min, each oligonucleotide was partially cleaved, generating a characteristic pattern of small bands. However, this activity was not dependent on the presence of the cA_4_ activator and therefore could not be associated with the activated toxin protein (Supplemental Figure 10B). Using the longest RNA substrate (RNA D), we observed that the Lon protease alone partly degraded the RNA after 60 min incubation at 60 °C, which may reflect a low level of nuclease contamination (Supplemental Figure 10C). In the presence of toxin, the same banding pattern observed in Supplemental Figure S10B was noted. We conclude that the activated toxin does not display specific cleavage activity against any of the five RNAs tested. The most likely explanation is that the toxin is specific for particular nucleic acid species or -structures.

## Discussion

The CRISPR-Lon protein is an unconventional example of a Lon protease, dedicated to specifically cut a protein and thereby releasing a MazF-like toxin. With a few exceptions such as viral proteases ^32, 34^, Lon proteases form large homo-hexameric ring-shaped complexes that processively unfold and degrade their targets in an ATP-dependent fashion. They are conserved from prokaryotes to eukaryotes and mitochondria and play essential roles in cell homeostasis, protein quality control, and metabolic regulation ^43–45^. CRISPR-Lon integrates the Lon protease domain into a completely different structural scaffold. While CRISPR-Lon is also activated by an ATP analog, it is not a processive enzyme.

CRISPR-Lon is the first type III-associated effector protein to be studied biochemically with a SAVED-instead of a CARF domain. The structural similarity with the cA_4_ binding CARF domains of the CRISPR endonuclease Can1 is striking, particularly given that the sequence similarity is very low. For the latter protein, an integrative approach involving small angle X-ray scattering revealed that cA_4_-binding induces large-scale conformational rearrangements of the effector domains that are thought to enable its nuclease activity ^19^. It is an interesting structural conundrum how binding of a cOA molecule to the sensor domains can activate effector domains that are connected in entirely different structural ways (Supplemental Figure 5), leading to the question of which mechanism might activate the CRISPR-Lon protease domain upon cA_4_ binding? For ATP-dependent Lon proteases, a recent study revealed that a conformational transition of an aspartic acid residue located three residues upstream of the catalytic serine is key to the activation mechanism ^46^. In the autoinhibited state of such proteases, the side chain forms a salt bridge with the lysine side chain of the catalytic dyad. In CRISPR-Lon, the corresponding residue is a serine, suggesting that the mechanism of activation/autoinhibition is different. Superpositions with other Lon proteases reveal that the helix (□J) containing K193 of the catalytic dyad is noticeably shifted, leading to a different relative orientation of the catalytic dyad residues (Supplemental Figure 4B). One might speculate that cA_4_ binding leads to a structural rearrangement of CRISPR-Lon or even of the whole CRISPR-Lon/T complex that enables the protease activity, releasing the CRISPR-T_23_ toxin-like domain. Our efforts to elucidate the function of CRISPR-T_23_ led to the conclusion that it is not an unspecific RNase, suggesting that it cleaves specific rRNA, mRNA or tRNA sequences as observed for other MazF toxins ^47–49^. As thought for other cOA dependent effectors, the activity of CRISPR-T might thus help to induce a dormant cell state, avoiding viral propagation and enabling the organism to have enough time to react against the attack ^7, 8^. If this was indeed true, the preformed CRISPR-Lon/T complex would be reminiscent of a “cocked gun” that could achieve a fast response mechanism, eliminating any delays associated with an upregulation of the toxin combined with the diffusion-limited activation of CRISPR-T by CRISPR-Lon. All organisms harboring CRISPR-Lon on their genomes have the CRISPR-T gene as neighbor. In the vicinity, all genes for CRISPR adaptation (Cas1-2) and interference (type III effector complex of subtype A, B, or D including Cas10) have been detected but no other CARF proteins such as the RNases Csm6 ^15^ or Csx1 ^8^. The release of a MazF RNase under control of the cOA-activated CRISPR-Lon protease would thus explain how the type III systems in those organisms operate (Figure 4).

**Figure 4:**
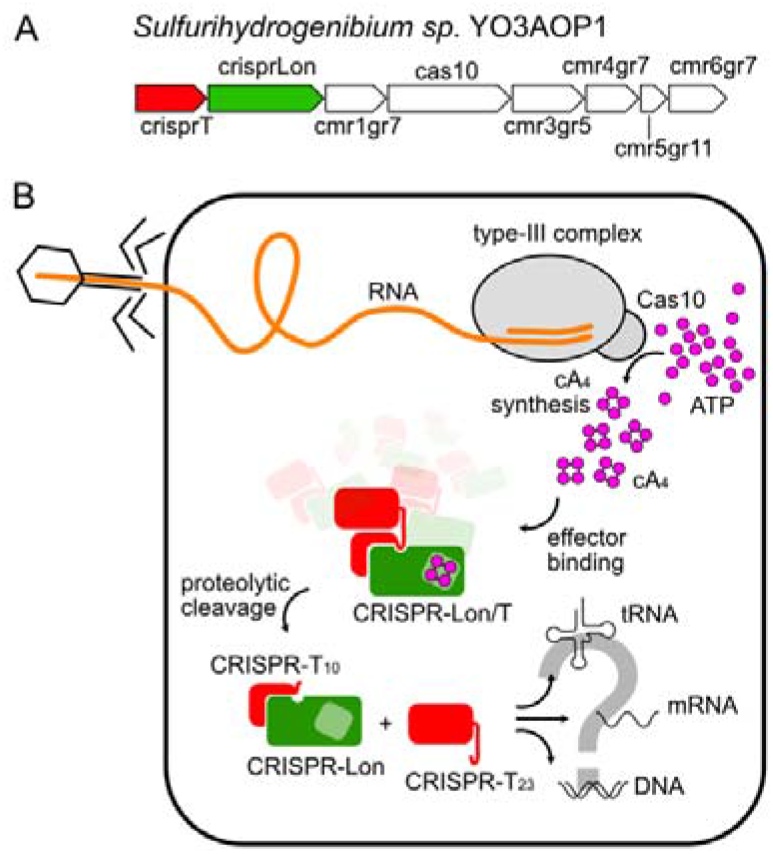
Model for CRISPR-Lon/T mediated antiviral defense. **A)** The *crisprLon* and *crisprT* genes are located in close proximity to the type III-B CRISPR genes of *Sulfurihydrogenibium sp.* YO3AOP1 (modified from ^9^). **B)** Once activated, the Cas10 subunit of the RNP synthesizes cA_4_ from ATP. The second messenger binds to the pool of preformed CRISPR-Lon/T complexes, which are quickly cleaved, releasing the MazF-like CRISPR-T_23_ fragments. The cellular target of the CRISPR-T_23_ fragment remains unclear but unspecific cleavage of RNA was not observed.

The use of proteases to trigger defense mechanisms is a well-known scheme in evolution. Prokaryotic type-II toxin-antitoxin (TA) systems, for instance, are activated by degradation of the antitoxin by ATP-dependent Lon proteases ^50^. While CRISPR-Lon/T combines the Lon protease fold and its activation by an ATP derivative (cA_4_) in a completely different structural arrangement, the functional scheme appears to be very similar. On the other end of the tree of life, the innate immune system of higher organisms also employs proteases such as caspases to initiate and amplify fast responses to external threats. Notably, caspase-like peptidases (“Craspases”) have also been discovered in connection to type III-D CRISPR systems, although the target of these enzymes is yet undefined ^51^.

Our study provides an interesting glimpse at both the possibilities and current limitations of structure prediction algorithms such as deepmind/alphafold2 ^40^. CRISPR-Lon was wrongly predicted as a transmembrane protein by algorithms that where state-of-the-art just a short while ago. In contrast, the deepmind/alphafold2 algorithm predicted the structure with astonishing accuracy (Supplemental Figure 2). However, a lot of functionally important information that comes with a high-resolution experimental structure, such as the position of ordered solvent molecules, is missing from the prediction. The second predicted structure, CRISPR-T, is not yet experimentally determined, but the prediction fits very nicely to our biochemical data of protease cleavage. In contrast, the CRISPR-Lon/T complex could not be predicted. Thus, structures of the CRISPR-Lon/T complex, the CRISPR-Lon/cA_4_ complex, and the question how the cA_4_ ligand activates the protease remain interesting and challenging tasks for the classical methods in structural biology.

## Methods

### Expression and purification of CRISPR-Lon

The codon-optimized gene for CRISPR–Lon was cloned into a pET11a vector with an N-terminal 10xHis-TEV tag. Site-directed mutagenesis was performed according to a protocol by Liu et al. ^52^. All CRISPR-Lon constructs were expressed in lysogeny broth (LB) medium.. *E. coli* BL21(DE3) cells were grown at 37° C until an OD_600_ of 0.6-0.8 was reached. Then, protein expression was started by induction with 0.4 mM IPTG, and the cell suspension was incubated at 30 °C for 4.5 h with shaking. Cells were harvested by centrifugation at 4,000*rcf for 25 min. at 20 °C and resuspended in lysis buffer (20 mM Tris, 50 mM NaCl, pH 8.0). The cells were lysed with a sonicator and cell debris was removed by centrifugation at 48,000*rcf for 45 min. at 4 °C. For protein purification, Ni^2+^-affinity chromatography (20 mM Tris, 50 mM NaCl, pH 8.0; 500 mM imidazole was included for elution) was followed by size-exclusion chromatography (20 mM Tris, 50 mM NaCl, pH 8.0) using a Superdex 200 16/600 column. After that, the His-tag was cleaved off by overnight incubation at 4 °C with a 1:50 molar ratio of protein to TEV protease (20 mM Tris, 50 mM NaCl, pH 8.0). A second Ni^2+^-affinity chromatography was used to remove the TEV protease and uncleaved protein. The purity of the protein was checked by SDS-PAGE after each purification step. After successful purification, the proteins were concentrated, flash-frozen in liquid nitrogen, and stored at - 80 °C in 20 mM Tris, 50 mM NaCl, pH 8.0. The selenomethionine derivative of CRISPR-Lon was prepared using *E. coli* B834 cells and the “SelenoMethionine Medium Complete” kit from Molecular Dimensions according to the instructions. Protein expression and purification were done in the same way as for the native protein.

### Expression and purification of CRISPR-T

The codon-optimized synthetic gene (BioCat) for CRISPR-T (UNIPROT-ID: B2V8L8), including an N-terminal 10xHis-TEV tag was cloned into a pET11a vector. Protein expression was done using the same expression strain and the same conditions as for CRISPR-Lon. Cells were harvested by centrifugation at 4,000*rcf for 25 min. at 20 °C and resuspended in lysis buffer (25 mM Tris, 500 mM NaCl, 10 % glycerol, 1 mM DTT, pH 8.0). The cells were lysed with a sonicator and cell debris was removed by centrifugation at 48,000*rcf for 45 min. at 20 °C. For protein purification, Ni^2+^-affinity chromatography (25 mM Tris, 500 mM NaCl, 1 mM DTT, 10 % glycerol, pH 8.0; 1 M imidazole was included for elution) was followed by size-exclusion chromatography (25 mM Tris, 500 mM NaCl, 1 mM DTT, 10 % glycerol, pH 8.0) using a Superdex 75 16/600 column. After that, the His-tag was cleaved off by overnight incubation at 4 °C with a 1:50 molar ratio of protein to TEV protease (25 mM Tris, 500 mM NaCl, 1 mM DTT, 10 % glycerol, pH 8.0). A second Ni^2+^-affinity chromatography was used to separate the TEV protease and uncleaved protein. The purity of the protein was checked by SDS-PAGE after each purification step. After successful purification, the proteins were concentrated, flash-frozen in liquid nitrogen, and stored at - 80 °C in 25 mM Tris, 500 mM NaCl, 1 mM DTT, 10 % glycerol, pH 8.0.

### Protease assay

For protease activity assays CRISPR–Lon and CRISPR–T were used at a final concentration of c = 4.64 µM each. The different cOAs were used at a final concentration of c = 5.11 µM. The protein solutions were prepared in 20 mM Tris, 50 mM NaCl, pH 8.0 and incubated for 60 min at 60 °C. Subsequently, the cOA was added and the mixture was incubated for another 60 min at 60 °C. For SDS-PAGE 3 µl of 4x SDS-loading buffer was added to 9 µl of the sample, the mixture was heated for 5 min at 94 °C and 10 µl were loaded to a 15 % polyacrylamide gel which was run at 250 V for 40 min.

### SEC-MALS

For determination of interactions between CRISPR–Lon and CRISPR–T, SEC-MALS runs were performed on an Agilent 1260 Infinity II Prime Bio LC System coupled with a Wyatt miniDAWN^®^ MALS detector using a Superose6 increase 10/300 chromatography column (GE Healthcare) equilibrated with 25 mM Tris, 500 mM NaCl, 1 mM DTT, 10 % glycerol, pH 8.0. Data acquisition and evaluation were carried out using ASTRA 8 software (Wyatt Technologies). The flow rate was set to 0.5 ml^*^min^1-^ and an injection volume of 50 µl was used for the experiments. Final concentrations were set to c(CRISPR–Lon) = 51 µmol*l^-1^, c(CRISPR–T) = 51 µmol*l^-1^ and c(cA4) = 60 µmol*l^1-^ by dilution with 25 mM Tris, 500 mM NaCl, 1 mM DTT, 10 % glycerol, pH 8.0.

### Mass spectrometry

Gel bands were cut from SDS-PAGE gels and sent to the Mass Spectrometry and Proteomics Facility of the University of St Andrews for analysis (https://mass-spec.wp.st-andrews.ac.uk).

### Ribonuclease assay

Ribonuclease activity of cleaved CRISPR-T (23 kDa fragment) was assayed by incubating full-length CRISPR-T with CRISPR-Lon and five different fluorescent-labelled RNA substrates, which were synthesised with the fluorescent dye (6-FAM) attached at 5’ end (purchased from Integrated DNA Technologies (IDT), Supplemental Table 2). The mixture of CRISPR-Lon (5.5 µM) and CRISPR-T (5.0 µM) was incubated at 60 °C in 20 mM Tris-HCl, pH 8.0, 50 mM NaCl and 0.5 mM EDTA for 30 min, cA_4_ (7.6 µM) was then added and the mixture was incubated for another 30 min at 60 °C, followed by adding one of the above RNA substrates into the mixture, incubating for an additional 60 min at 60 °C. Finally, 6 µl of the sample was analysed on SDS-PAGE (NuPAGE Bis-Tris Gel, Thermo Fisher Scientific) by heating at 95 °C for 5 min with 2 µL of SDS-PAGE loading dye (Thermo Fisher Scientific; μ NuPAGE Sample Reducing Agent and LDS Sample Buffer). The remaining 14 µl of the sample were loaded to 20 % acrylamide, 7 M urea, 1×TBE denaturing gel, which was run at 30 W, 45 °C for 2 h. The gel was finally imaged by Typhoon FLA 7000 imager (GE Healthcare) at a wavelength of 532 nm (pmt 600∼700).

### X-ray crystallography

Pure CRISPR-Lon protein was concentrated to 20 mg/ml and crystallized using a Gryphon pipetting robot (Art Robbins) and commercial crystallization screens (Molecular Dimensions). Hexagonal crystals appeared after one day in condition D7 of the JCSG+ screen. Several rounds of optimization were performed to achieve well-diffracting crystals. The final crystallization condition was 0.1 M Tris-Cl pH 8.0, 38.8 % PEG 400, 0.29 M Li_2_SO_4_. The SeMet derivative (see above) was crystallized under similar conditions and yielded identical crystals. The crystals were harvested without further cryo-protection and a diffraction dataset was recorded at beamline P13 operated by EMBL Hamburg at the PETRA III storage ring (DESY, Hamburg, Germany) ^53^. The diffraction data were automatically processed with XDS ^54^. The structure was solved using phenix.autosolve and refined with phenix.refine ^55^. Model building was performed in Coot ^56^ and figures were prepared with PyMOL (www.pymol.org). The geometry of the model was checked with MolProbity ^57^.

### Structural predictions with deepmind/alphafold2

The source code of the deepmind/alphafold2 algorithm was downloaded from https://github.com/deepmind/alphafold and installed as described https://github.com/deepmind/alphafold. The algorithm was run using the CASP14 preset. The pLDDTs scores were extracted and mapped onto the structures with PyMOL. The CRISPR-Lon/T complex was predicted by concatenating the CRISPR-T and CRISPR-Lon amino acid sequencing separated by a 30 amino acid linker consisting of “U” residues (“unkown”).

### Data availability statement

The crystal structure has been deposited in the PDB (XXXX).

### Code availability statement

No custom code was used in this work.

## Acknowledgements

The synchrotron MX data were collected at beamline P13, operated by EMBL Hamburg at the PETRA III storage ring (DESY, Hamburg, Germany). We would like to thank Gleb Bourenkov and Isabel Bento for the assistance in using the beamline. We thank Virginius Siksnys for helpful discussions. We thank Sally Shirran for the mass spectrometry analysis. We would like to thank Norbert Brenner for technical assistance.

## Author contributions

C.R. and G.H. conceived and supervised the study and performed initial protein expression and crystallization experiments. N.S., C.R., and G.H. designed experiments. N.S. optimized the purification of CRISPR-Lon and CRISPR-T, crystallized CRISPR-Lon and established and performed the cleavage assays and SEC-MALS experiments. M.F.P. cloned the cleavage site mutants. G.H. and N.S. solved and refined the CRISPR-Lon structure. H.C. and M.F.W. planned and performed the ribonuclease assay. C.R., M.F.W., H.C., N.S., and G.H. analyzed the data and wrote the paper. R.S. and W.B. cloned the initial CRISPR-Lon construct. All authors discussed the results and commented on the manuscript at all stages.

---

**Supplemental Table 1:** Data collection and refinement statistics.

|  |  |
| --- | --- |
| Wavelength (Å) | 0.9795 |
| Resolution range (Å) | 54.06 - 2.1 (2.175 - 2.1) |
| Space group | P 3<sub>2</sub>21 |
| Unit cell (Å, °) | 68.90, 68.90, 255.38, 90, 90, 120 |
| Total reflections | 83686 (7464) |
| Unique reflections | 41843 (3732) |
| Multiplicity | 2.0 (2.0) |
| Completeness (%) | 99.02 (90.28) |
| I/sigma(I) | 23.20 (2.36) |
| Wilson B-factor (Å<sup>2</sup>) | 36.42 |
| R-merge | 0.054 (0.3444) |
| R-meas | 0.07637 (0.4871) |
| R-pim | 0.054 (0.3444) |
| CC1/2 | 0.995 (0.705) |
| CC* | 0.999 (0.909) |
| Reflections used in Refinement | 41842 (3732) |
| Reflections used for R-free | 1996 (175) |
| R-work | 0.1932 (0.2489) |
| R-free | 0.2255 (0.3097) |
| CC(work) | 0.962 (0.818) |
| CC(free) | 0.941 (0.741) |
| Number of non-hydrogen atoms | 4487 |
| Ramachandran outliers/favored (%) | (0/97) |
| Clashscore | 4.53 |
| Molprobity score | 1.43 |
| Macromolecules | 4175 |
| Ligands | 69 |
| Solvent | 243 |

Numbers in parenthesis refer to the shell of highest resolution.

**Supplemental Table 2:**
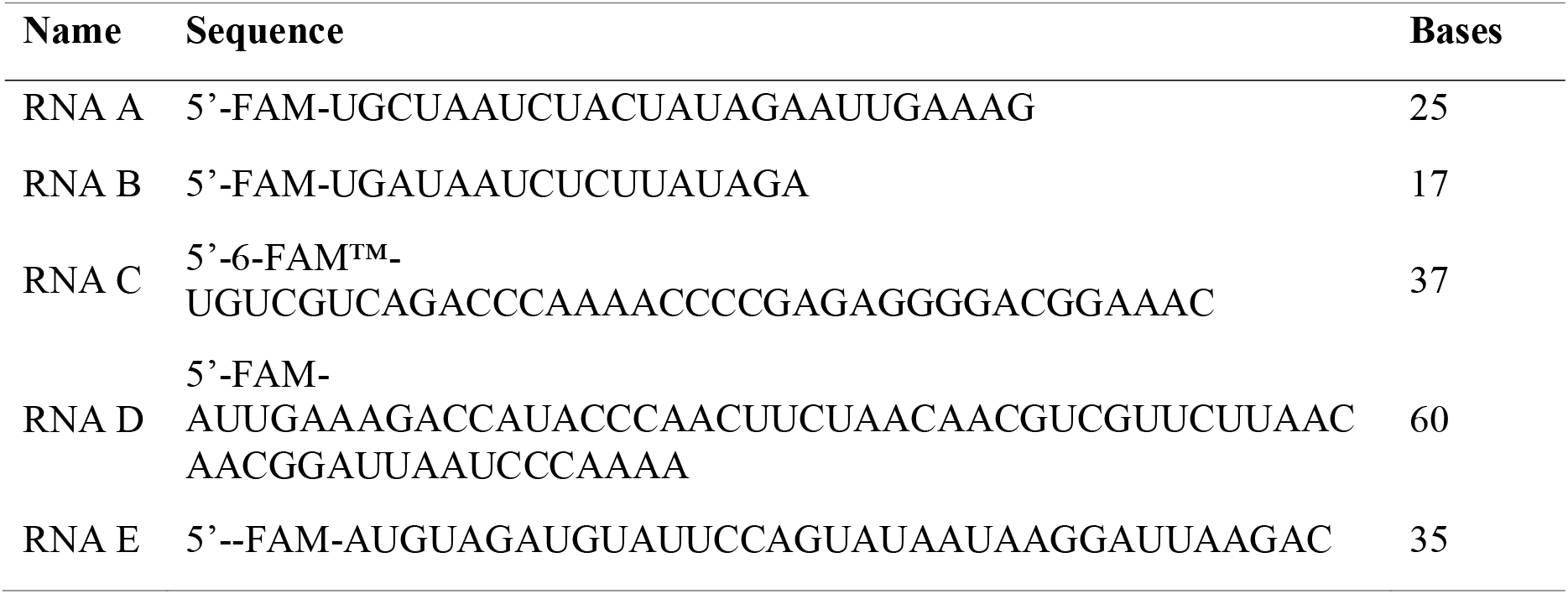
Substrate for the ribonuclease assay.

**Supplemental Figure 1:**
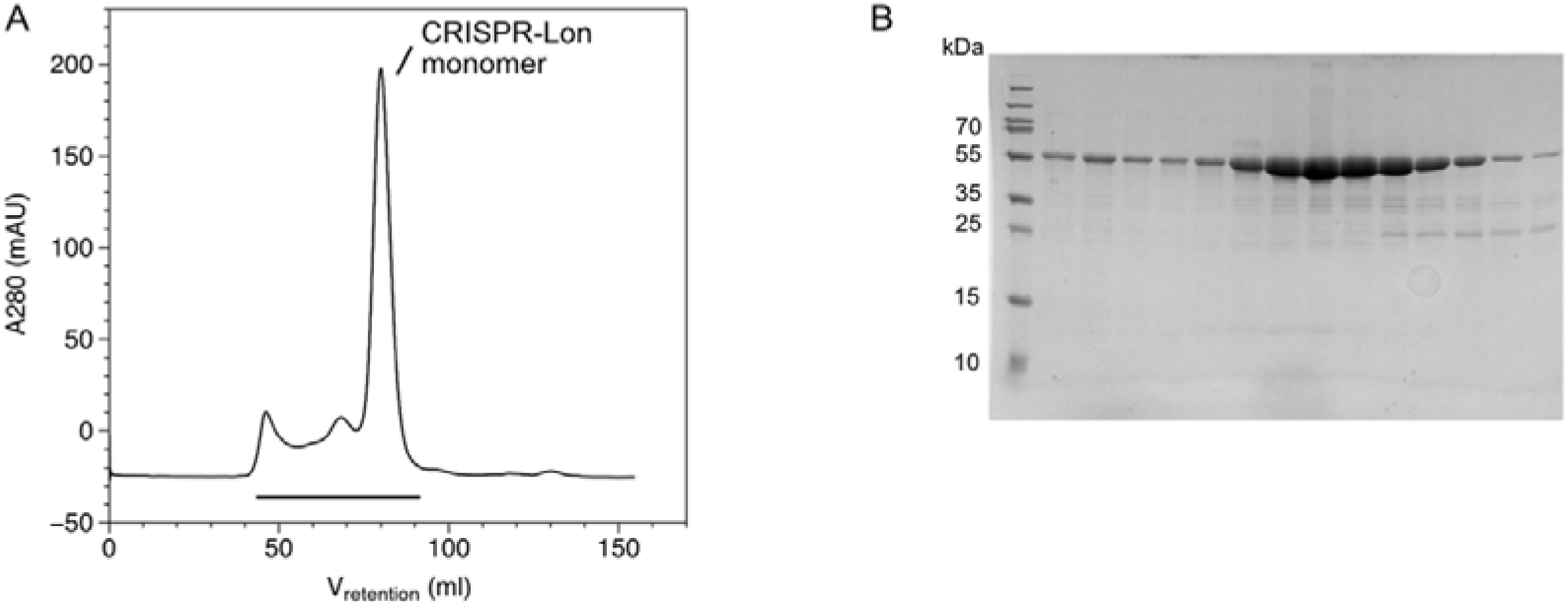
Purification of CRISPR-Lon. **A)** Gelfiltration chromatography (Superdex 200 16/60) of CRISPR-Lon. The protein elutes as a monomer. **B)** SDS-PAGE analysis of the fractions indicated by the black bar in A).

**Supplemental Figure 2:**
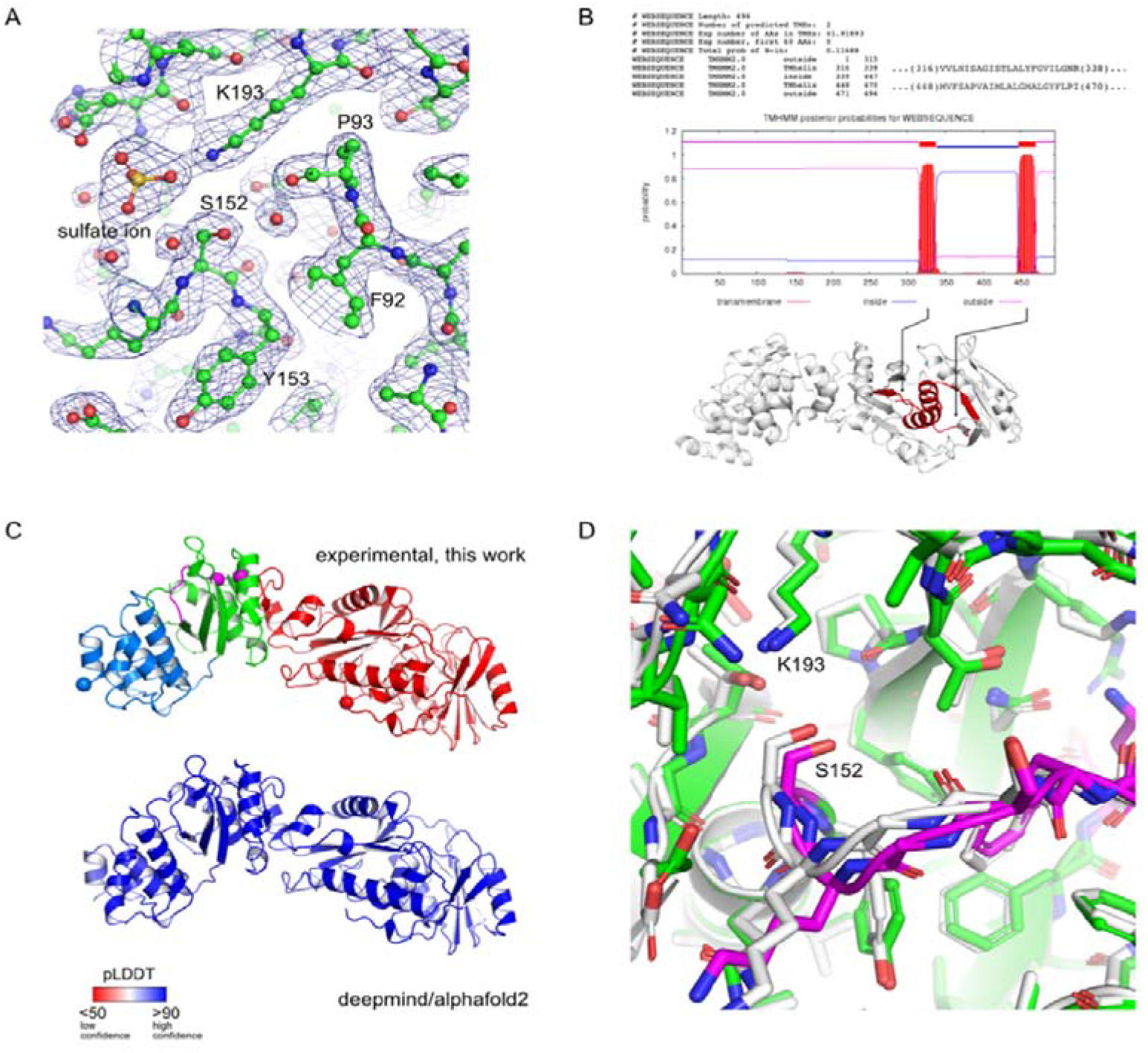
Refinement and model building. **A)** Representative electron density of the SeMet CRISPR-Lon crystal structure. The structural model is shown in ball-and-stick representation. Selected residues are labeled. The blue mesh is a 2mFo-DFc electron density map contoured at 1.0 σ. **B)** TM-prediction by the TMHMM 2.0 server^35^ vs experiment. **C)** Deepmind/alphafold2 vs experiment. **D)** Close-up of the Lon protease active site region with the catalytic dyad residues S152 and K193 labelled.

**Supplemental Figure 3:**
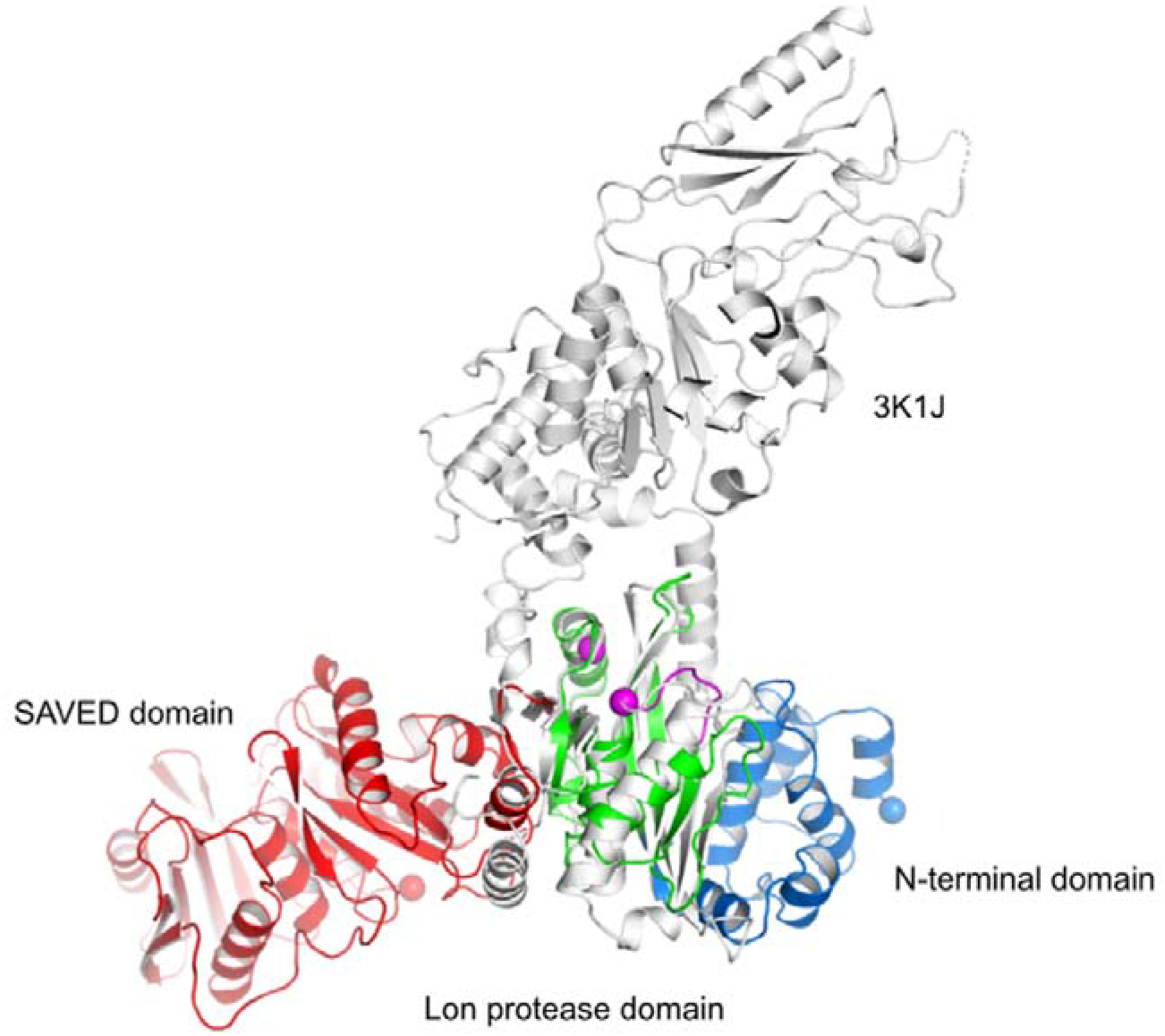
Superposition of CRISPR-Lon active site with the ATP-dependent Lon protease from *Thermocuccus onnorineus*. CRISPR-Lon is shown as a cartoon model color-coded as in Figure 1. The Lon protease from *T. onnorineus* (PDB-ID: 3K1J, DALI Z-score: 12.8, ^31^) is shown as a white cartoon model.

**Supplemental Figure 4:**
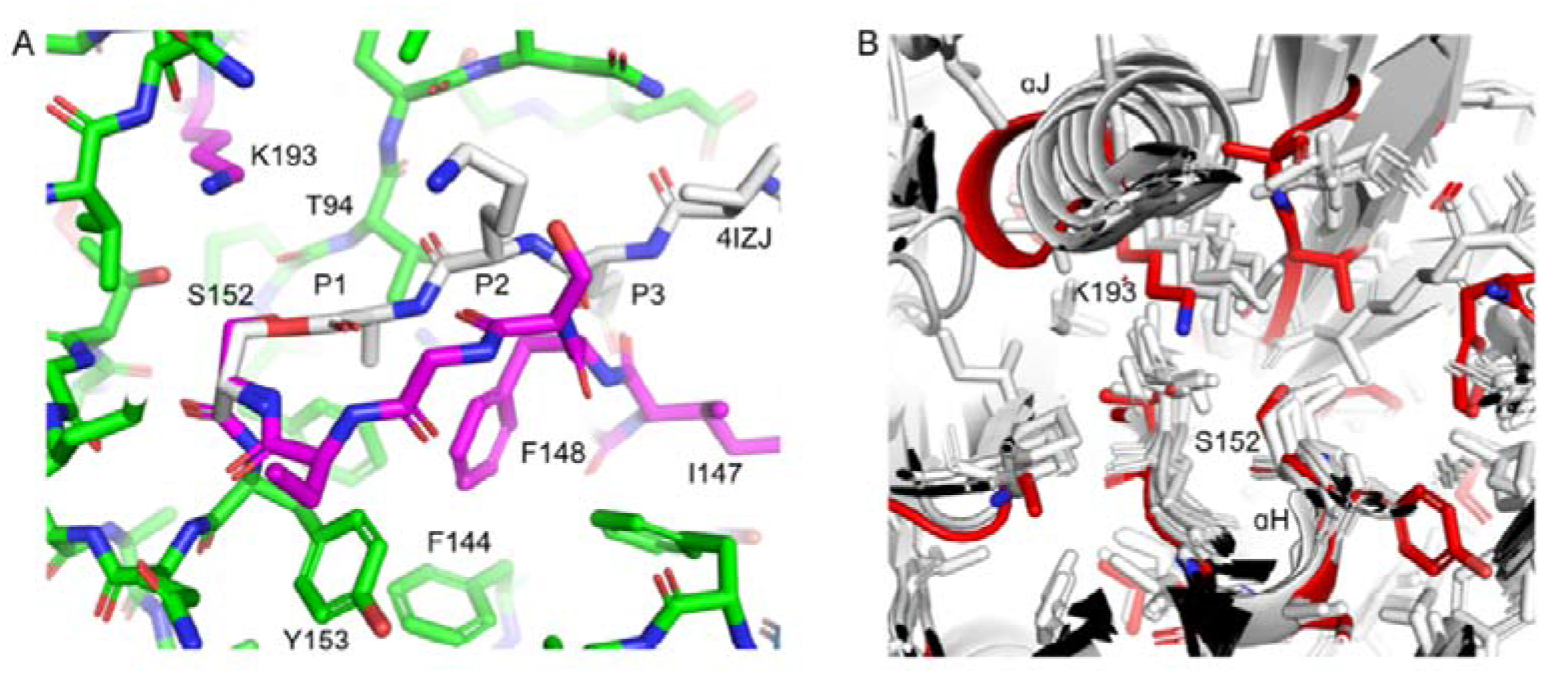
A) Superposition of CRISPR-Lon active site with the acyl-enzyme intermediate of yellowfin asciitis virus protease. **A)** CRISPR-Lon is shown in sticks representation and color-coded as in Figure 1. Chain D of structure 4IZJ ^34^ (residues 630-640) was superimposed on the corresponding residues of CRISPR-Lon (150-160) leading to an r. m. s. d. of 0.314 Å. Of 4IZJ, only the acyl-enzyme intermediate is shown in sticks mode. Selected residues and the positions of the P1-P3 sites are indicated. **B)** Comparison of the CRISPR-Lon (red) protease active site with related structures (white). The structures were superposed based on the catalytic serines and the □-helices immediately downstream of it (□H in CRISPR-Lon). The catalytic dyad residues of CRISPR-Lon are marked.

**Supplemental Figure 5:**
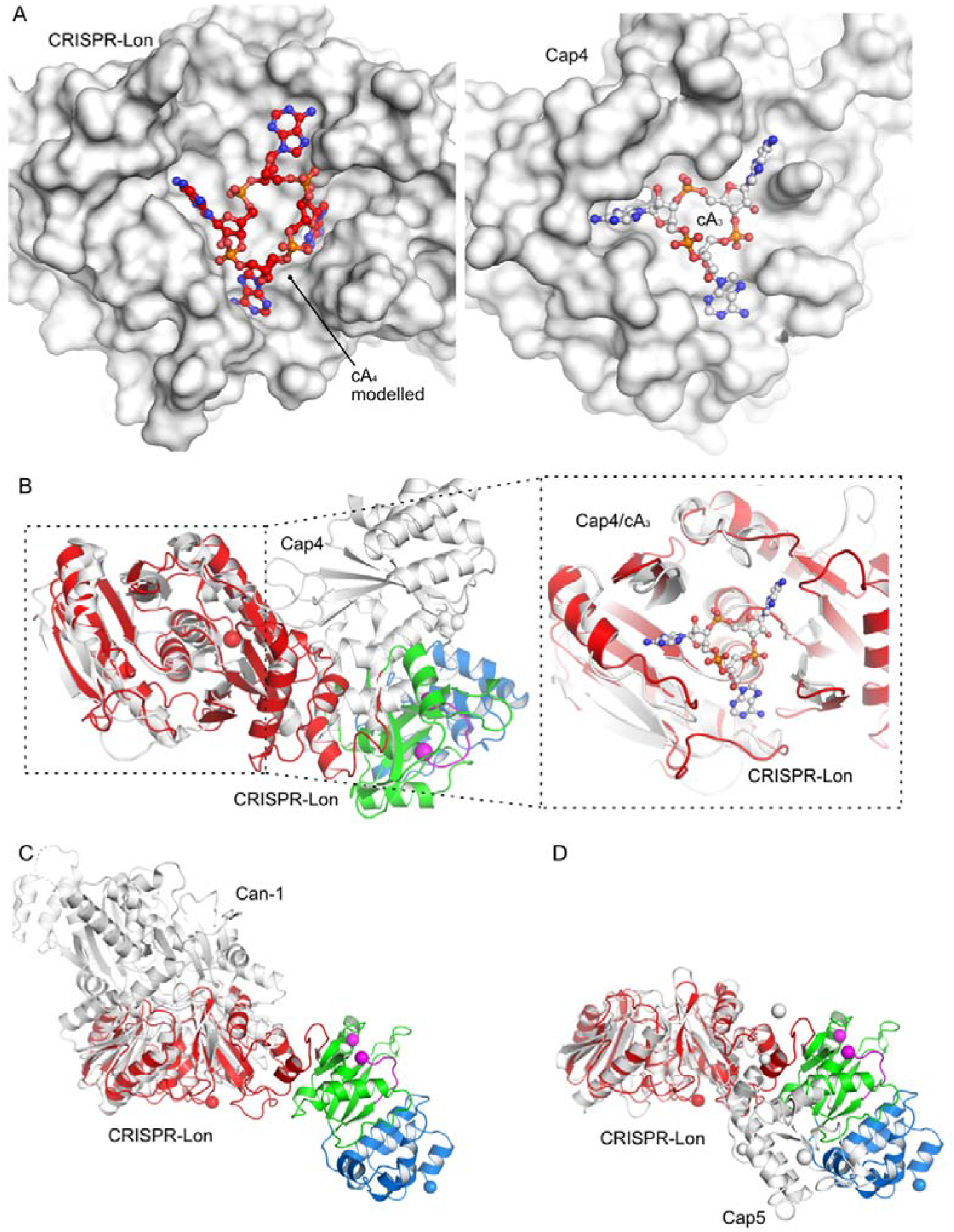
The SAVED domain of CRISPR-Lon. **A** Left: A cA_4_ molecule (red balls-and-sticks) was modeled into the cOA binding site of the CRISPR-Lon SAVED domain (white surface). Right: The Cap4/cA_3_ complex structure. The cA_3_ molecule is shown as white balls-and-sticks and Cap4 as a white surface (PDB-ID: 6VM6 ^24^). Both CRISPR-Lon and Cap4 are oriented as in B). **B)** Left: Superposition of CRISPR-Lon (color scheme as in Figure 1) with the Cap4 protein (white, PDB-ID: 6VM6 ^24^). Right: Close-up of the superposed SAVED domains. The cA_3_ molecule of the Cap4 complex structure is shown as white ball-and-stick model (PDB-ID: 6WAN ^24^). **C)** Superposition of the CRISPR-Lon SAVED domain (color scheme as in Figure 1) with the CARF domains of the Can1 protein (white, PDB-ID: 6SCE ^19^). **D)** Superposition of the CRISPR-Lon SAVED domain (color scheme as in Figure 1) with the Cap5 protein (white, PDB-ID: 7RWK ^36^).

**Supplemental Figure 6:**
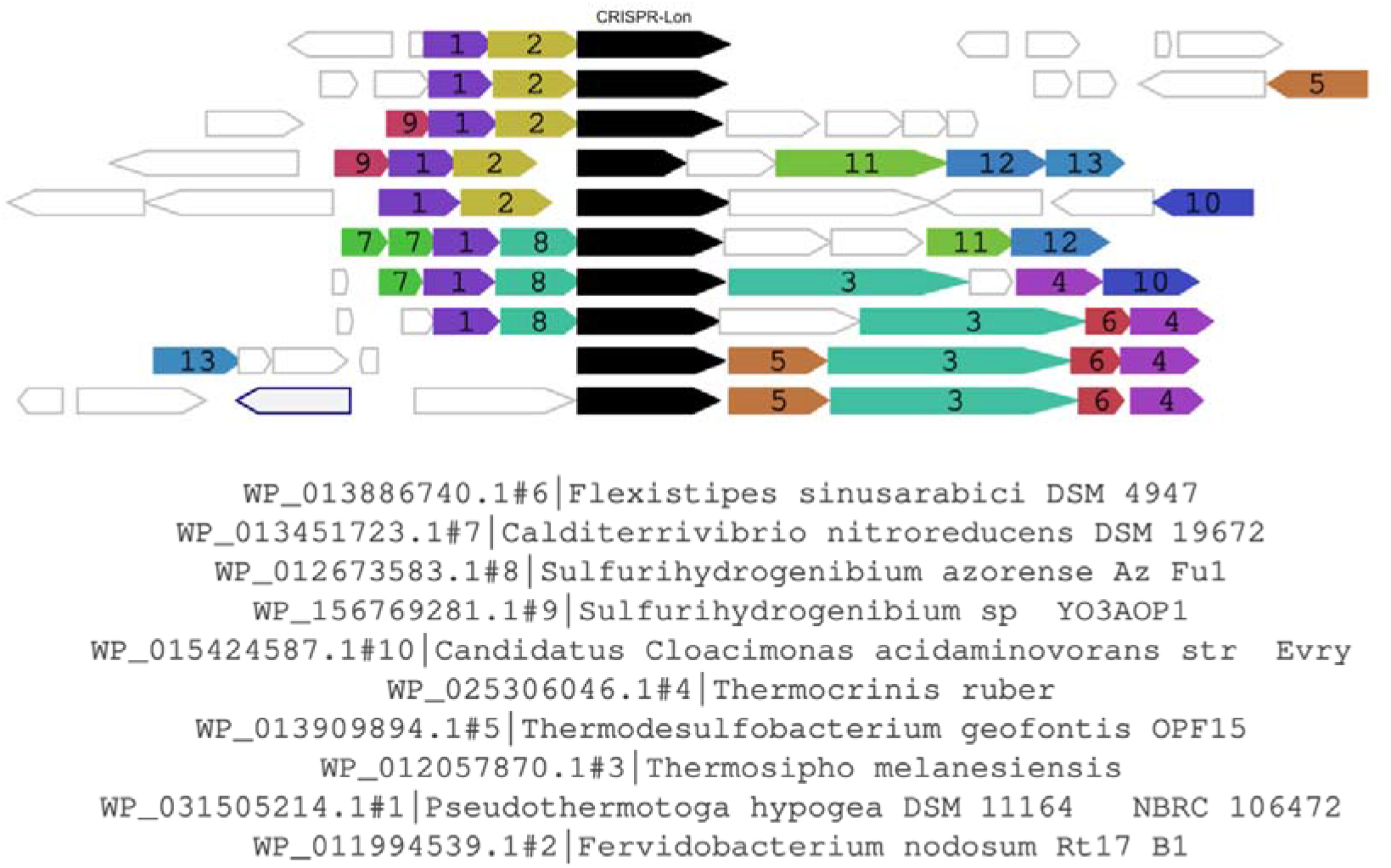
Genetic neighborhood of CRISRP-Lon as determined by the WebFLAGs^38^ server. The CRISPR-Lon gene is shown as a black arrow and the CRISPR-T gene as a yellow arrow. Genes of a certain color have similar sequences that form numbered groups. Genes without filling color are not conserved in the given context. Grey genes with blue borders are pseudogenes.

**Supplemental Figure 7:**
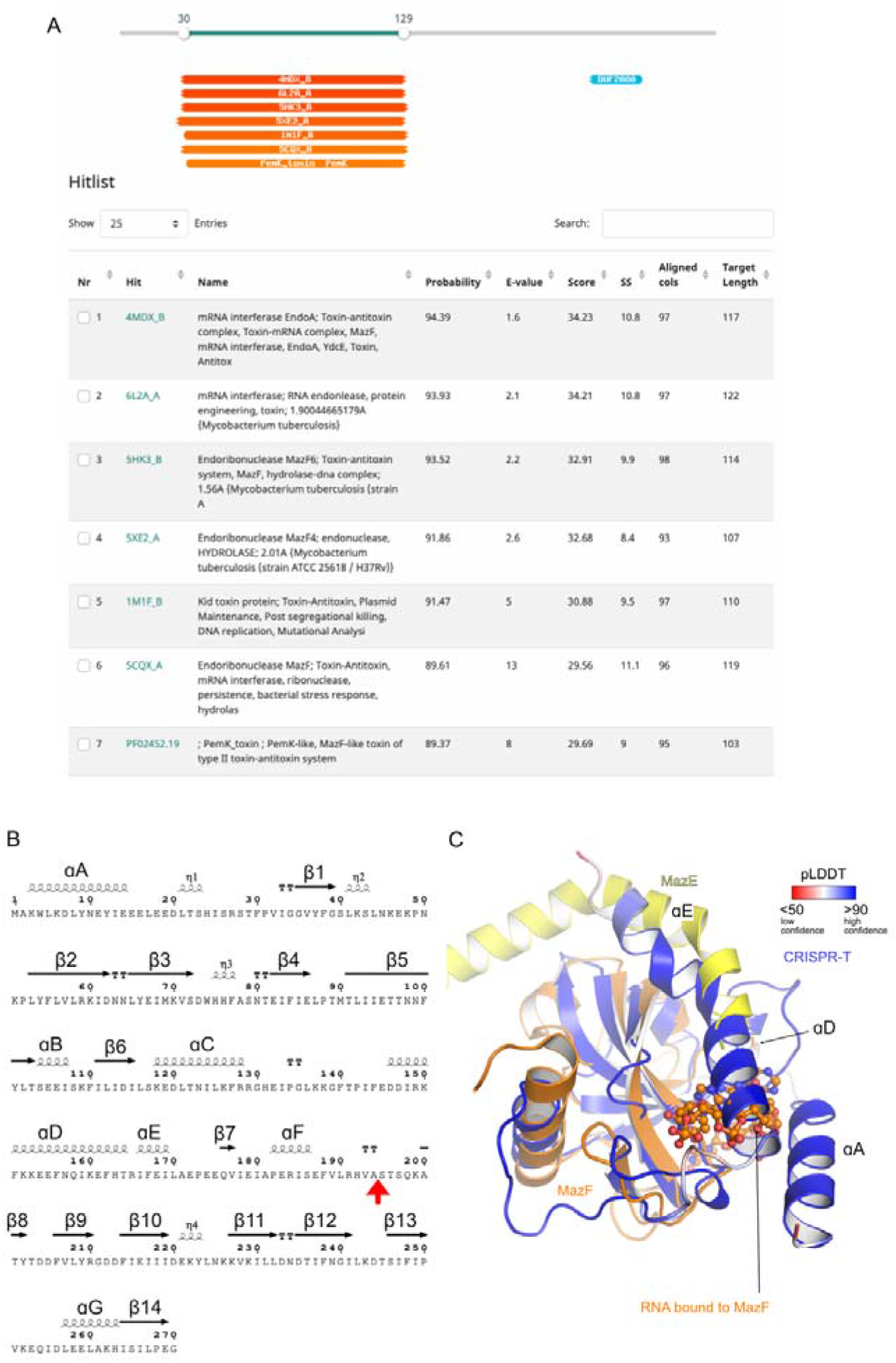
Bioinformatic information about CRISPR-T. **A)** HHPred ^39^ result for CRISPR-T. **B**) Secondary structure of the predicted CRISPR-T (Figure 2B) structure mapped to its amino acid sequence (ESPRIPT ^58^). The CRISPR-Lon cleavage site is marked with a red arrow. **C)** A superposition of the predicted CRISPR-T structure (compare Figure 2B) with the MazE/F (orange/yellow) complex and the MazF/ssRNA complex (only the RNA is shown as spheres model) (PDB-IDs: 4ME7 ^59^ and 5CR2 ^48^). The deepmind/alphafold2 ^40^ prediction confidence is mapped onto the CRISPR-T structure. Note that the MazE/F complex is dimeric and only one half of the full complex is shown for clarity.

**Supplemental Figure 8:**
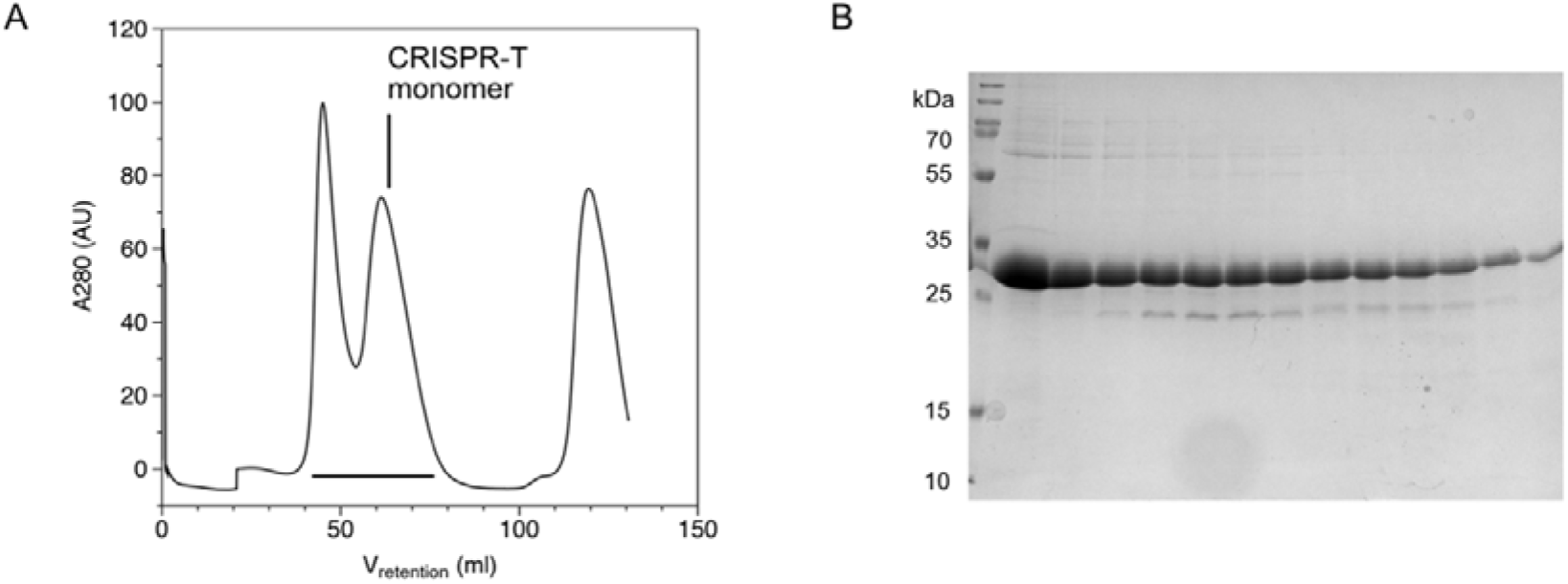
Purification of CRISPR-T. **A)** Gelfiltration chromatography (Superdex 75 16/60) of CRISPR-T. The protein elutes as a monomer. **B)** SDS-PAGE analysis of the fractions indicated by the black bar in A).

**Supplemental Figure 9:**
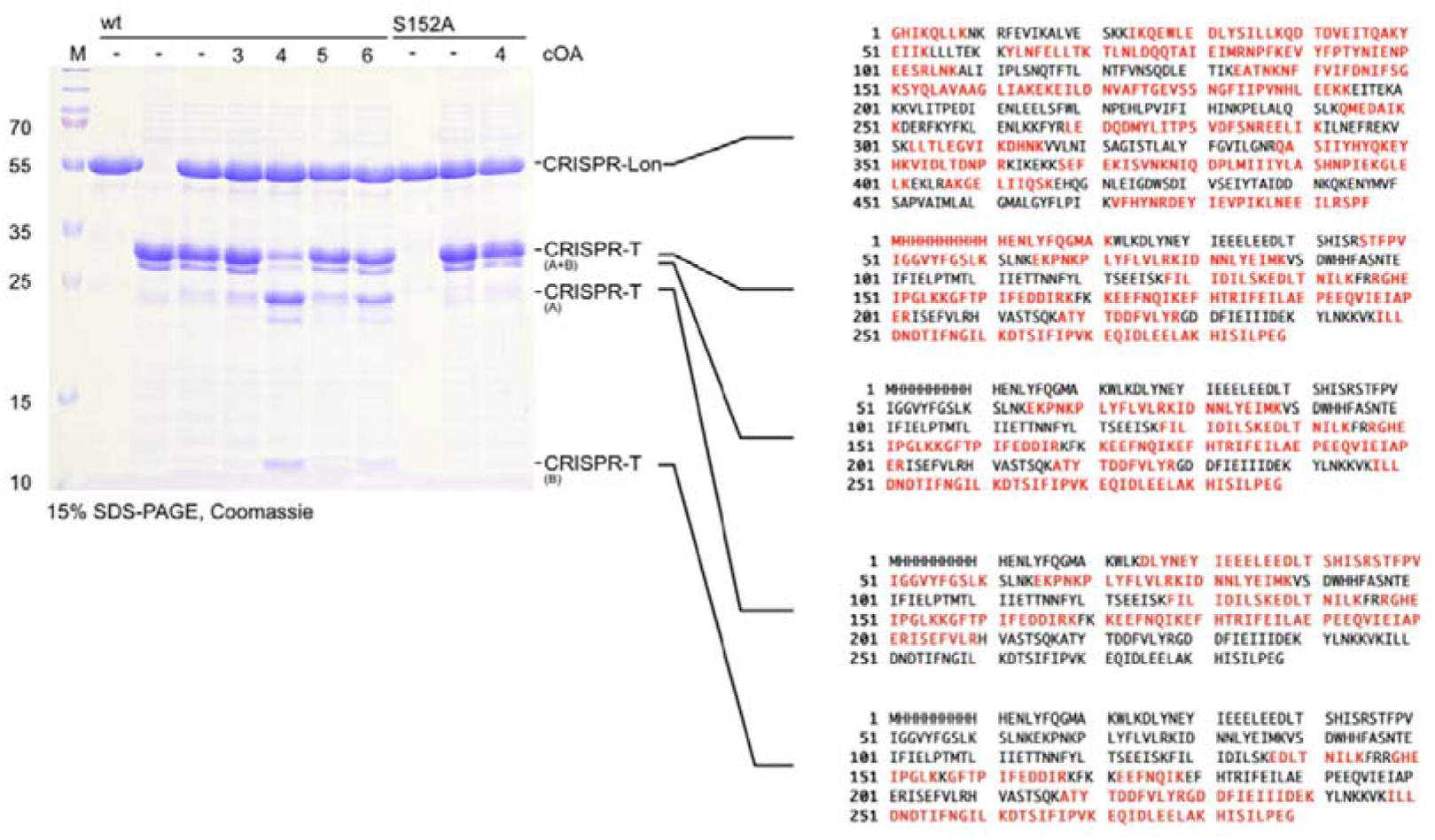
Peptide fingerprints of cleavage bands. The indicated gel-bands were cut from the gel and submitted for identification at the Mass spectrometry and proteomics facility at the University of St Andrews (Fife, UK, https://mass-spec.wp.st-andrews.ac.uk). Red letters indicate peptides that were identified in the respective sample.

**Supplemental Figure 10:**
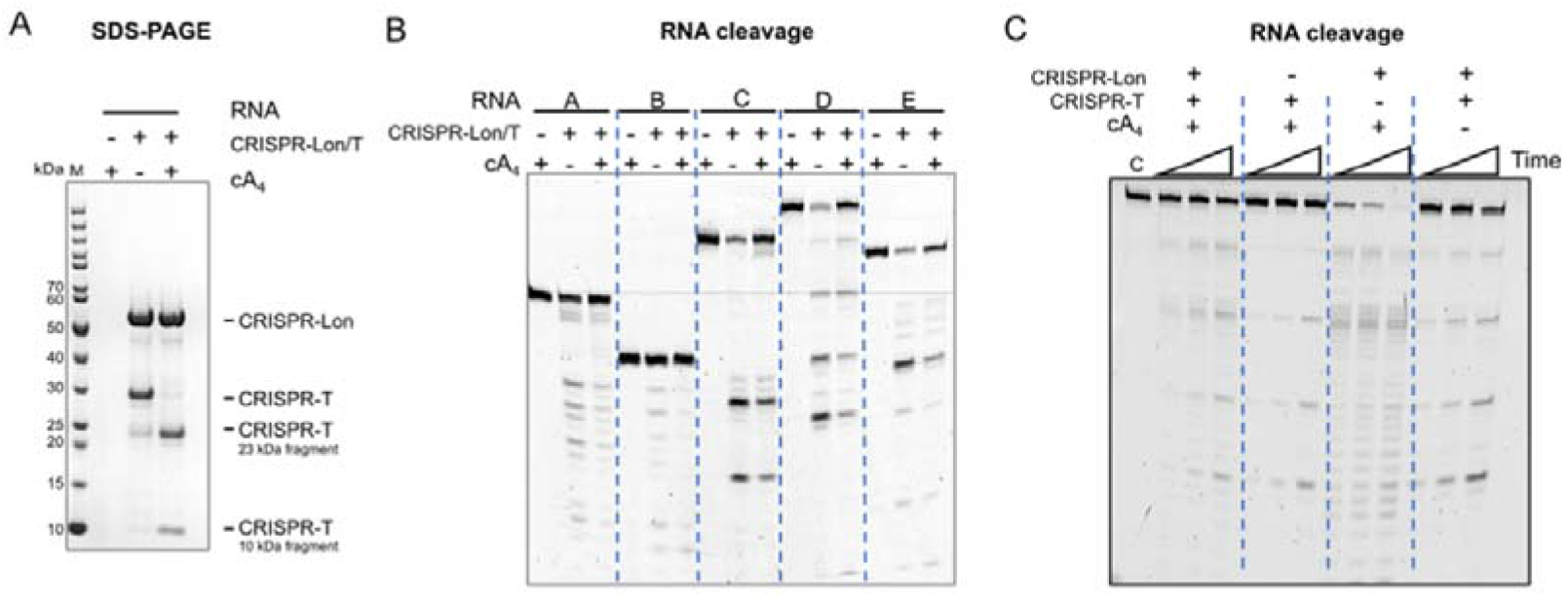
Probing the RNase activity of the activated toxin. **A)** SDS-PAGE analysis of cA_4_-induced cleavage of CRISPR-T (33 kDa) by CRISPR-Lon. Cleavage is complete after 60 min at 60 °C. **B)** Fluorescence image of the denaturing polyacrylamide gel electrophoresis to determine ribonuclease activity of the reactions shown in A) against five fluorescently-labelled RNA substrates (RNAs listed in Supplemental Table 2 correspond to set A, B, C, D and E). Some cleavage reactions were observed after 60 min incubation at 60 °C, but these were not dependent on the presence of cA_4_ activator. **C)** Activity of CRISPR-Lon and CRISPR-T toward fluorescent-labelled RNA substrate D. Reaction mixture was incubating for 5, 10 and 30 min at 60 °C. Control reaction only contains the RNA substrate. The Lon protease degraded the RNA substrate, suggesting the presence of an RNase contaminant. The same banding pattern seen in B was observed when CRISPR-T was present, but this was not dependent on the presence of the protease or the cA_4_ activator.

